# A deeply conserved miR-1 dependent regulon supports muscle cell physiology

**DOI:** 10.1101/2020.08.31.275644

**Authors:** Paula Gutiérrez-Pérez, Emilio M. Santillán, Thomas Lendl, Anna Schrempf, Thomas L. Steinacker, Mila Asparuhova, Marlene Brandstetter, David Haselbach, Luisa Cochella

## Abstract

Muscles are not only essential for force generation but are also key regulators of systemic energy homeostasis^1^. Both these roles rely heavily on mitochondria and lysosome function as providers of energy and building blocks, but also as metabolic sensors^2-4^. Perturbations in these organelles or their crosstalk lead to a wide range of pathologies^5^. Here, we uncover a deeply conserved regulon of mitochondria and lysosome homeostasis under control of the muscle-specific microRNA miR-1. Animals lacking miR-1 display a diverse range of muscle cell defects that have been attributed to numerous different targets^6^. Guided by the striking conservation of miR-1 and some of its predicted targets, we identified a set of direct targets that can explain the pleiotropic function of miR-1. miR-1-mediated repression of multiple subunits of the vacuolar ATPase (V-ATPase) complex, a key player in the acidification of internal compartments and a hub for metabolic signaling^7,8^, and of DCT-1/BNIP3, a mitochondrial protein involved in mitophagy and apoptosis^9,10^, accounts for the function of this miRNA in *C. elegans*. Surprisingly, although multiple V-ATPase subunits are upregulated in the absence of miR-1, this causes a loss-of-function of V-ATPase due to altered levels or stoichiometry, which negatively impact complex assembly. Finally, we demonstrate the conservation of the functional relationship between miR-1 and the V-ATPase complex in *Drosophila*.

---

miR-1 is one of only 32 miRNAs that are conserved throughout Bilateria^11,12^. Its conservation extends from its sequence to its muscle-specific expression, and to its profound impact on muscle development and physiology in every species studied. Loss of miR-1 causes defects in autophagy, sarcomere structure and mitochondrial integrity, thus affecting the energy homeostasis and general physiology of muscle cells, and impairing myoblast differentiation and survival^6,13-21^. In contrast to most miRNAs, target prediction indicates that miR-1 targets are also conserved. Strikingly, miR-1 has predicted binding sites in transcripts encoding many of the 15 subunits that make up the V-ATPase complex in worms, flies, fish, mice and humans^22-24^ (**Extended Data Fig. 1a,b**). Whereas deregulation of some V-ATPase subunits has been observed upon loss of miR-1^18^, no functional link between miR-1 and the V-ATPase has been established – although the central role of the V-ATPase in lysosomal degradation and metabolic signaling^7,8^ makes it a prime candidate to account for the broad function of miR-1.

In addition to the V-ATPase, *dct-1/bnip3* and *tbc-7/ tbc1d15* also have predicted miR-1 binding sites in their 3′ UTRs across multiple animals (and/or sites for miR-133, another muscle-specific miRNA clustered with miR-1) (**Extended Data Fig. 1a**). DCT-1/BNIP3 is a key regulator of mitophagy and, under stress conditions or aging, it is critical to preserve mitochondria turnover and cellular homeostasis^9,10,25^. TBC-7/TBC1D15 is a Rab GTPase-activating protein that regulates lysosomal morphology and fusion between late endosomes and lysosomes^26^. It has recently been shown that loss of miR-1 in *C. elegans* and mammalian cells causes TBC-7/TBC1D15 overexpression, leading to impaired autophagy and proteotoxic stress^20^. Given the involvement of the V-ATPase, DCT-1/BNIP3 and TBC-7/TBC1D15 in proteostasis, autophagy and metabolism, we hypothesized that they could constitute a conserved regulon that accounts for the diverse range of cellular defects observed in miR-1 mutant muscles.

To test this, we took advantage of *C. elegans* genetics and the fact that while loss of miR-1 causes lethality in other animals^13-17^ miR-1-deficient worms are viable in the lab, albeit with a number of muscle-cell defects^19,20^. To further characterize the muscles of miR-1-deficient *C. elegans*, we first visualized the sarcomeric structure of the body-wall muscle by electron microscopy, immunostaining and live-imaging of transgenic animals carrying a myosin heavy chain (MYO-3) fused to GFP, using two different mutant alleles of *mir-1* (**Extended Data Fig. 2a**). In contrast to other animals^15,16,18^, we observed no obvious loss of sarcomere integrity (**Extended Data Fig. 2b,c**). However, similar to mouse skeletal muscle, where loss of miR-1 causes disruption of mitochondrial structure and function^21^, we found that miR-1-deficient L1 larvae displayed significant fragmentation of the otherwise extensive tubular mitochondrial network in the body-wall muscle (**Fig. 1a, Extended Data Fig. 2d**). Accompanying the morphological defect, we measured reduced ATP levels in the body-wall muscle as measured with a fluorescent ATP sensor^27^ (**Fig. 1b**). Moreover, using an aggregation prone polyglutamine-YFP expressed in the cytoplasm of body-wall muscles^28^, we observed an increase in number of aggregates in the absence of miR-1, this has also recently been reported in an independent study^20^ (**Fig. 1c**).

**Fig. 1.**
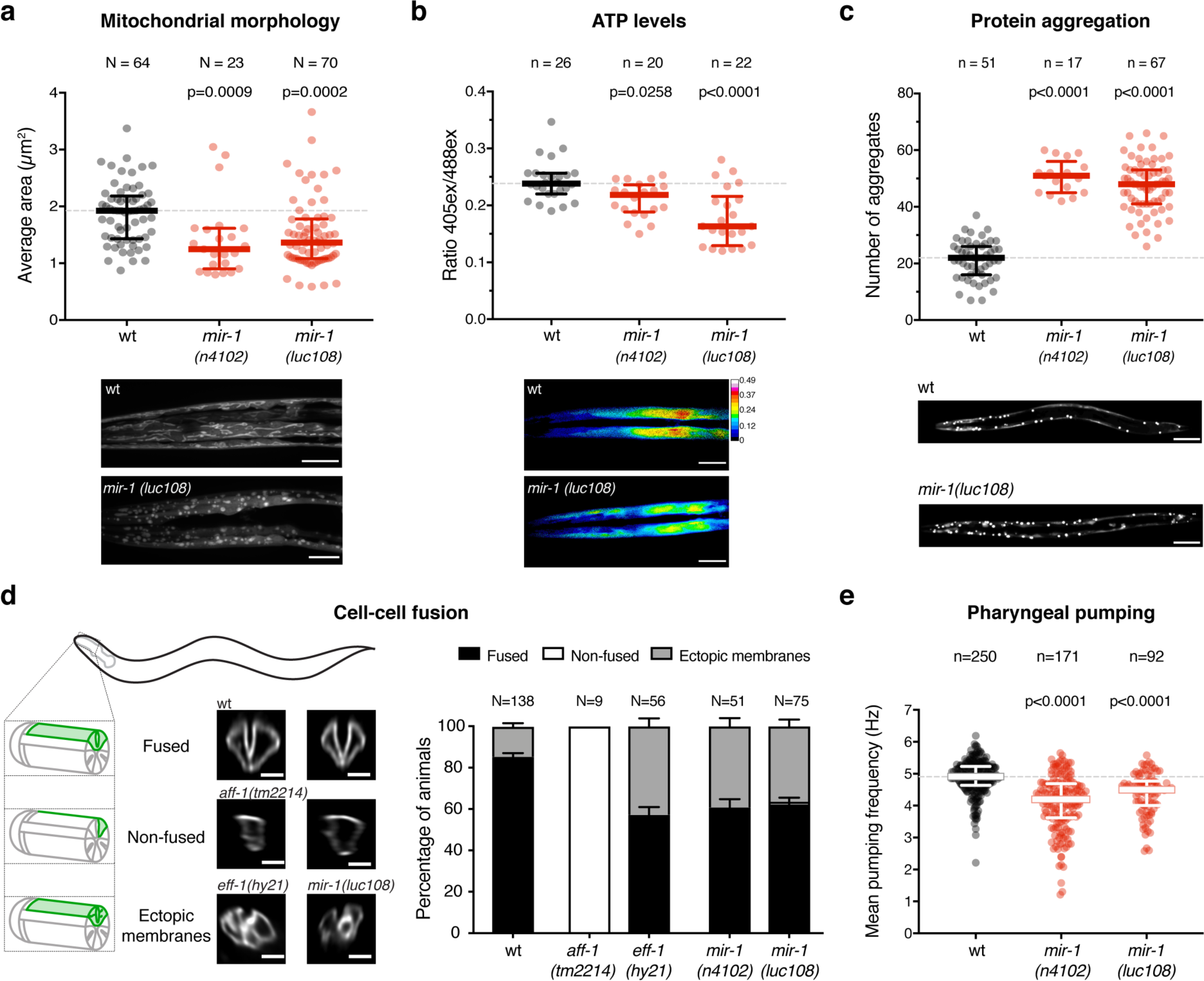
Absence of miR-1 causes diverse muscle defects in *C. elegans*. **a**. Live L1 larvae of the indicated genotypes, expressing a mitochondria-localized GFP reporter in the body-wall muscle, were imaged using confocal microscopy. Automated analysis (see Methods) was used to extract average mitochondria area (among other parameters, **Extended Data Fig. 2d**). *mir-1(0)* animals have smaller mitochondria in muscle. **b**. Live L1 larvae expressing a ratiometric ATP sensor^27^ were imaged using confocal microscopy. Automated analysis (see Methods) was used to extract the emission ratio between 405 and 488 excitation wavelengths. *mir-1(0)* animals have reduced ATP concentration in muscle. **a,b**. P-values by Kruskal-Wallis test are indicated. Representative images are shown; scale bars = 10µm. **c**. L4 larvae of the indicated genotypes, expressing the aggregation-prone Q40::YFP reporter^28^ were imaged and the number of aggregates per animal was counted. *mir-1(0)* animals have more aggregates. P-values by one-way ANOVA are indicated. Representative images are shown; scale bars = 50µm. **d**. (left) Schematic of the cell-cell fusion assay: pm3DL and R fusion is assessed by the expression of a pm3DR-specific membrane marker (green). If fusion occurs, the whole pm3D outline is labelled (fused), otherwise only pm3DR remains labelled (non-fused). Membrane trafficking or fusion defects may result in ectopic membranes. Representative images are shown. Scale bars = 20µm. (right) Quantification of the cell-cell fusion events. **e**. Pumping frequencies of wt and *mir-1(0)* animals were extracted from electropharyngeograms (EPGs) conducted in Nemametrix ScreenChips in the presence of 10 mM serotonin. *mir-1(0)* animals have reduced pumping frequency. P-values by Kruskal-Wallis test are indicated. N/n refers to number of animals analyzed.

In mammals, loss of miR-1 results in decreased cell-cell fusion, impairing myotube formation^29,30^. While body-wall muscles do not fuse in *C. elegans*, pharyngeal muscle cells do^31^; we thus developed an assay to monitor the fusion of two specific pharyngeal muscle cells, pm3DL and pm3DR (**Fig. 1d**). We monitored fusion in animals defective for miR-1 but also for the known cell-cell fusogens in *C. elegans*, EFF-1 and AFF-1^32^. pm3DL/R fusion was completely abolished in *aff-1-*mutants, as expected^33^. Animals deficient for *eff-1* or for miR-1 looked indistinguishable from each other but different from *aff-1*, displaying ectopic labeled membranes either internally, or forming bridges between the two muscle cells (**Fig. 1d**). This may be related to previously described defects in autofusion in *eff-1* mutants^34-36^ and suggests that miR-1-deficient animals have decreased expression or activity of EFF-1. Notably, trafficking and recycling of EFF-1 are altered upon V-ATPase loss-of-function in the *C. elegans* hypodermis^37,38^, providing a potential link.

Finally, we asked if these cellular defects translated into locomotion or pharyngeal pumping defects. We did not observe differences in posture or speed of locomotion between wild-type and miR-1-deficient L4/young adult animals (not shown), but we did measure a decrease in pharyngeal pumping frequency in miR-1 mutants (**Fig. 1e**). This was due to extended duration of each pumping cycle, specifically during the muscle relaxation phase (**Extended Data Fig. 2e,f**). Overall, *C. elegans* muscle cells lacking miR-1 display a broad range of cellular defects similar to those observed in other animals.

To assess if the defects observed in miR-1-deficient animals are due to the de-repression of V-ATPase subunits and/or of DCT-1/BNIP3, we used CRISPR/ Cas9 to mutate the predicted miR-1-binding sites in the 3′ UTRs of *dct-1* (*dct-1*^*NotI*^) or of six *vha* genes coding for V-ATPase subunits simultaneously (*6x-vha*^*NotI*^). The selected *vha* genes have conserved predicted miR-1 binding sites and constitute different parts of the complex (the V_0_ membrane ring, the V_1_ peripheral ring or part of the connector between both) (**Fig. 2a, Extended Data Fig. 1**). The levels of all six *vha* transcripts in the *6x-vha*^*NotI*^ mutant strain were increased relative to wild type and mirrored those in miR-1-deficient animals, indicating that miR-1 is a direct repressor of multiple *vha* genes (**Extended Data Fig. 3a,b**). The levels of *dct-1* and *tbc-7* transcripts were also increased in the absence of miR-1 (**Extended Data Fig. 3c**).

**Fig. 2.**
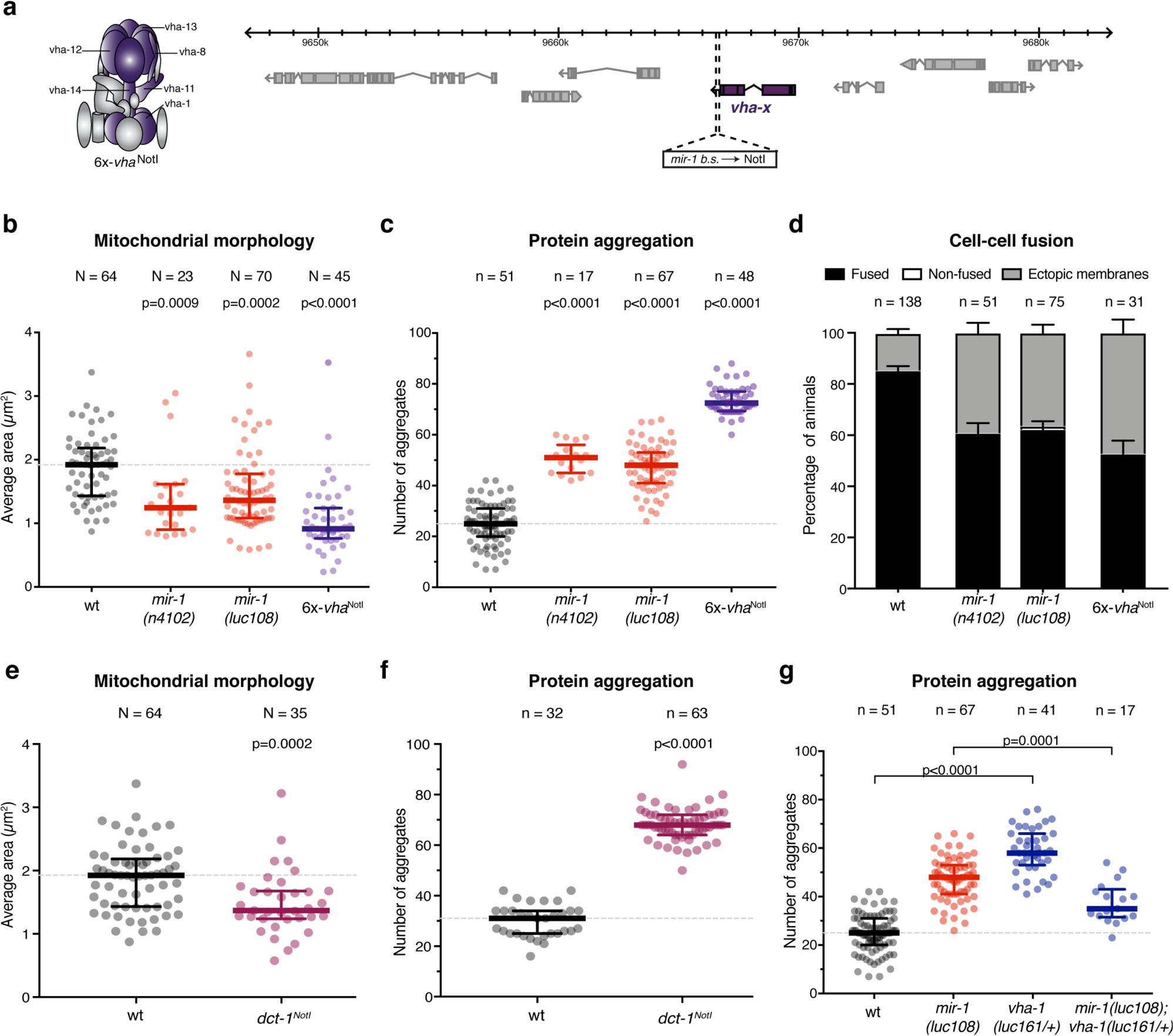
Direct repression of V-ATPase subunits and DCT-1/BNIP3 by miR-1 is necessary for muscle homeostasis. **a**. Schematic representation of the design of the *6x-vha*^*NotI*^ strain. The *mir-1* binding sites in the 3’UTRs of *vha-1, -8, -11, -12, -13* and *-14* (purple) were replaced by NotI restriction sites. **b**. Average mitochondria area, extracted by automated analysis of images as in Fig. 1a, is significantly reduced in *6x-vha*^*NotI*^ L1 larvae. P-values by Kruskal-Wallis test are indicated. **c**. Number of aggregates per animal measured as in Fig. 1c; *6x-vha*^*NotI*^ L4 animals show impaired proteostasis. P-values (one-way ANOVA) are shown. **d**. Quantification of the different fusion outcomes as in Fig. 1d; *6x-vha*^*NotI*^ L1 animals also display ectopic membranes in pharyngeal muscle cells. wt and *mir-1(0)* data sets are replotted from Fig. 1d. **e**. Average mitochondria area is significantly reduced in *dct-1*^*NotI*^ L1 larvae. P-value by Mann-Whitney test is indicated. **f**. *dct-1*^*NotI*^ L4 animals show an increased in *Q40::YFP* aggregates. P-value (unpaired t-test) is shown. **g**. Reduction in *vha-1* levels rescues *mir-1(0)* proteostasis defects, as indicated by a significant decrease in the number of Q40::YFP aggregates per animal. P-values (one-way ANOVA) are shown. wt and *mir-1(0)* datasets for both mitochondria and protein aggregation assays are replotted from Fig. 1a and c, respectively. N/n refers to number of animals analyzed.

If transcripts encoding V-ATPase subunits and/or DCT-1/BNIP3 are functionally relevant targets of miR-1, removal of the miR-1 binding sites from their 3′ UTRs should cause the same range of defects as loss of miR-1 itself. We thus assessed mitochondrial morphology, protein aggregation and pharyngeal muscle fusion in the *6x-vha*^*NotI*^ strain (**Fig. 2b-d**), and the mitochondria and proteostasis-related phenotypes in the *dct-1*^*NotI*^ strain (**Fig. 2e,f**). Both were in otherwise wild-type backgrounds in which miR-1 is present and can regulate all other putative targets. Remarkably, de-repression of the *vha* transcripts or of *dct-1*, was sufficient to cause the same defects we observed in *mir-1(0)* animals (**Fig. 2b-f**). Providing additional support for the functional connection between the V-ATPase and miR-1, we observed that removing one genomic copy of *vha-1* reduced the number of protein aggregates in *mir-1* mutant animals (**Fig. 2g**). These results indicate that the V-ATPase and DCT-1/BNIP3 are functional targets of miR-1 and also, that they act non-redundantly in a shared pathway or process such that deregulation of either one has a negative impact on protein and mitochondria homeostasis.

The mitochondrial fragmentation and protein aggregation caused by loss of miR-1 parallel the defects observed upon BNIP3 and TBC1D15 overexpression^9,10,20^, consistent with the expected upregulation of these proteins upon removal of a repressor. Conversely, increased mitochondrial fragmentation and protein aggregation have been linked to *decreased* activity of the V-ATPase complex, leading to impaired lysosomal degradation, autophagy and metabolic signaling^39-43^. This suggests that although multiple *vha* transcripts are upregulated in the absence of miR-1-mediated repression, this results in a loss-of-function of the complex. We hypothesized that de-repression of one or more subunits could disrupt the required levels or stoichiometry for the correct assembly of this intricate 15-subunit complex^44,45^, resulting in reduced V-ATPase function.

To test this, we first confirmed the effects on protein aggregation and mitochondrial fragmentation of partial loss-of-function of the V-ATPase complex (complete loss-of-function results in embryonic lethality). Deletion of subunits with redundant counterparts (*vha-2/3*) or heterozygous mutation of unique subunits (*vha-1, -5* and *-8*) caused significant increases in mitochondrial fragmentation and protein aggregation (**Fig. 2g, Fig. 3a,b**), phenocopying the miR-1 deletion.

**Fig. 3.**
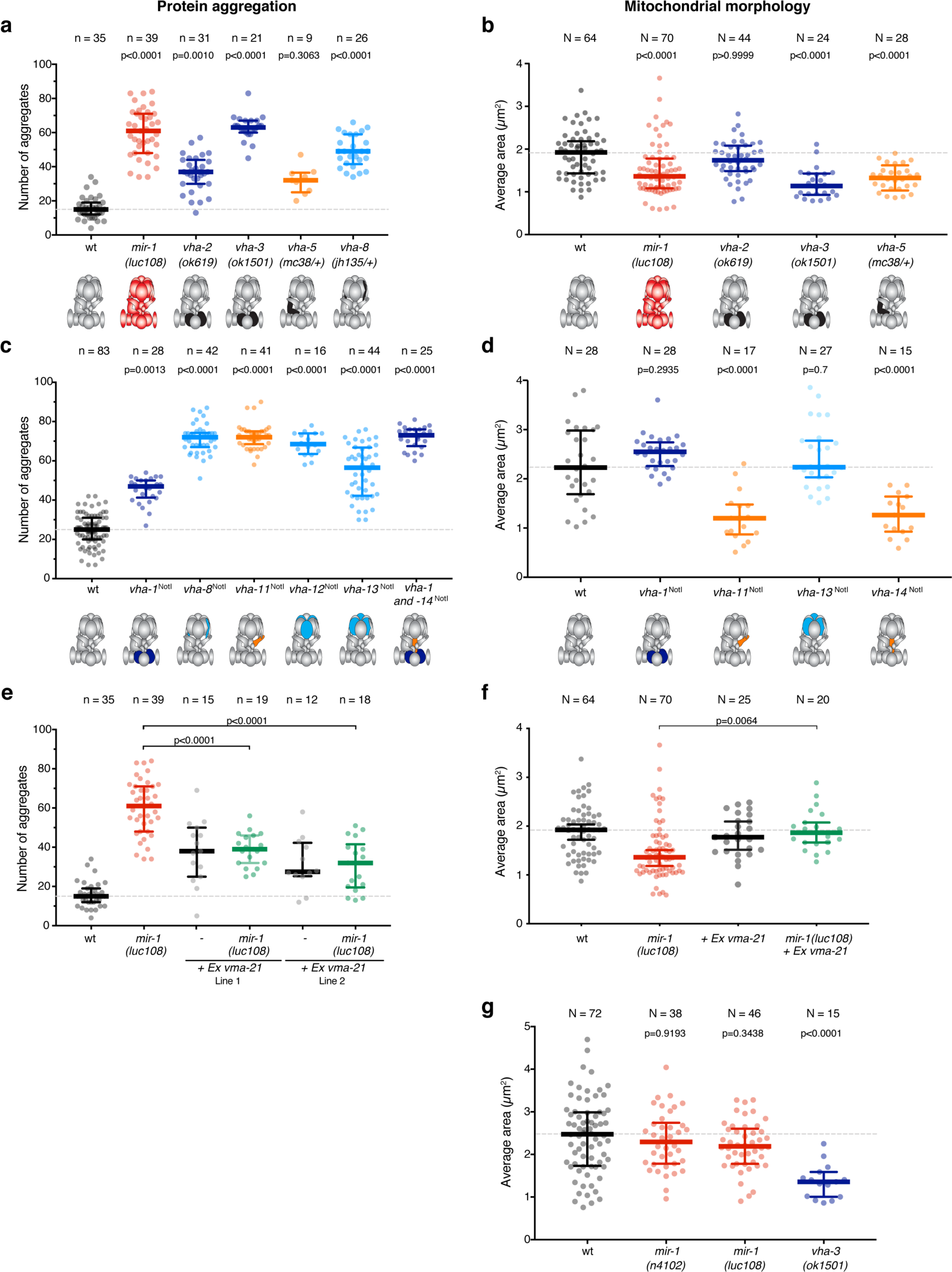
Upregulation of V-ATPase subunits in the absence of miR-1 results in decreased function of the V-ATPase complex due to assembly defects. **a** and **b**. V-ATPase loss of function mutations in *vha-2, -3, -5 or -8* results in increased protein aggregation (a) and mitochondria fragmentation (b). wt and *mir-1(0)* datasets in (a) are replotted from Fig. 1a. P-values by Kruskal-Wallis test are indicated. V-ATPase complex schematics are represented below, where black represents loss of function of each particular subunit. **c**. De-repression of single subunits is sufficient to cause proteostasis defects. P-values by Kruskal-Wallis test are indicated. V-ATPase complex schematics represent which subunit carries miR-1 binding site mutations and is thus de-repressed, dark blue represents a V0 membrane-integral subunit, orange a key subunit for assembly and light blue a V1 subunit involved in the hydrolysis of ATP. **d**. De-repression of *vha-11* and *vha-14* alone, but not of *vha-1* or *-13* is sufficient to cause mitochondrial network fragmentation. wt and *mir-1(0)* datasets are replotted from Fig. 1a. P-values by Kruskal-Wallis test are indicated. **e**. Overexpression of VMA-21 rescues miR-1 proteostasis defects. Two independent extrachromosomal lines are analyzed. P-values by one-way ANOVA are shown. wt and *mir-1(0)* datasets are replotted from Fig. 3b. **f**. Overexpression of VMA-21 rescues miR-1 mitochondrial network fragmentation. P-values by Kruskal-Wallis are shown. wt and *mir-1(0)* datasets are replotted from Fig. 1a. **g**. Starvation rescues mitochondrial network fragmentation in *mir-1(0)* animals but not in *vha-3(0)* mutants. P-values by one-way ANOVA are shown. N/n refers to number of animals analyzed.

We then tested how deregulation of individual V-ATPase subunits impacts mitochondrial structure and proteostasis. In the loss-of-function scenario, the upregulation of each individual subunit could create abnormal interactions affecting the assembly pathway of the complex and in turn resulting in decreased V-ATPase function. In the alternative “increased V-ATPase function” model, only the upregulation of one limiting subunit, or all limiting subunits simultaneously, would lead to formation of more V-ATPase complex. We found that removing the miR-1 binding sites from every tested subunit individually, caused a similar extent of protein aggregation as loss of miR-1 (**Fig. 3c**). Also, de-repression of *vha-11* or *vha-14* alone resulted in mitochondrial fragmentation (**Fig. 3d**). The fact that de-repression of various individual subunits is sufficient to cause these defects is difficult to reconcile with an increased function model and rather supports a model in which loss of miR-1 causes loss-of-function of the V-ATPase.

We reasoned that loss of V-ATPase function could arise from complex assembly defects. In particular, formation of the membrane-embedded V_0_ ring requires dedicated chaperones in the endoplasmic reticulum (ER)^45^. One of these chaperones, VMA21, is necessary in stoichiometric amounts to assist V_0_ assembly in the ER and its shuttling to the Golgi, where the V_0_ and the peripheral V_1_ subcomplex come together_46_. We thus asked whether overexpression of the homolog of VMA21 in *C. elegans*, R07E5.7^47^, could alleviate the defects observed in miR-1-deficient animals. Notably, overexpression of the ER chaperone VMA21 significantly decreased the number of cytoplasmic aggregates as well as the mitochondria fragmentation in animals carrying the *mir-1* deletion (**Fig. 3e,f**). These data suggest that both these defects stem from decreased V-ATPase function, at least in part due to impaired assembly of the V_0_ domain.

Glucose or amino acid deprivation has been shown to promote V-ATPase activation in mammalian cells by promoting association of the V_0_ and V_1_ subcomplexes^48,49^. We thus asked if starvation of L1 larvae had a beneficial effect on miR-1 mutant animals. Consistent with an improvement in V-ATPase function, the mitochondria morphology of starved miR-1-deficient animals was similar to the wild type (**Fig. 3g**). In contrast, starvation was unable to rescue mitochondria fragmentation in animals in which V-ATPase loss-of-function was due to genomic deletion of a subunit (**Fig. 3g**). Together, these observations suggest a role for miR-1 in maintaining different V-ATPase subunits at the right dose or stoichiometry to enable correct assembly of this crucial multi-subunit complex in muscle cells.

Finally, we set out to test whether the remarkable conservation of the miR-1 sequence and the presence of predicted miR-1 binding sites in multiple V-ATPase subunits also reflects a functional interaction in an arthropod, estimated to have diverged from nematodes ∼600 Mya. To this end, we analyzed *Drosophila melanogaster* wild-type and miR-1-deficient larvae. Flies deficient for miR-1 have various muscle defects which result in growth arrest as second instar larvae and lethality within 2-4 days of hatching^16,17^. We first quantified the levels of various transcripts encoding V-ATPase subunits in first instar larvae and found that multiple *Vhas* were upregulated in the absence of miR-1 (**Extended Data Fig. 3e**). To test the functional connection between miR-1 and the V-ATPase complex, we asked whether reducing the levels of one of the subunits involved in early stages of assembly might rescue the larval growth arrest. Akin to what we observed in *C. elegans*, deletion of one genomic copy of the V_0_ component *Vha100-3* partially rescued the growth defect of miR-1-deficient larvae (**Fig. 4a**). In contrast, reducing the level of one of the members of the peripheral V_1_ ring did not significantly rescue the growth defect (**Fig. 4a**). This is consistent with our observations in *C. elegans* pointing to loss of miR-1 causing a V-ATPase assembly defect, primarily of the V_0_ part of the complex. Concomitant with the increased growth, reducing the level of *Vha100-3* also significantly extended the survival of miR-1-deficient larvae (**Fig. 4b**). Together, these results strongly support the existence of a conserved functional interaction between miR-1 and the V-ATPase that deserves further exploration in other animals.

**Fig. 4.**
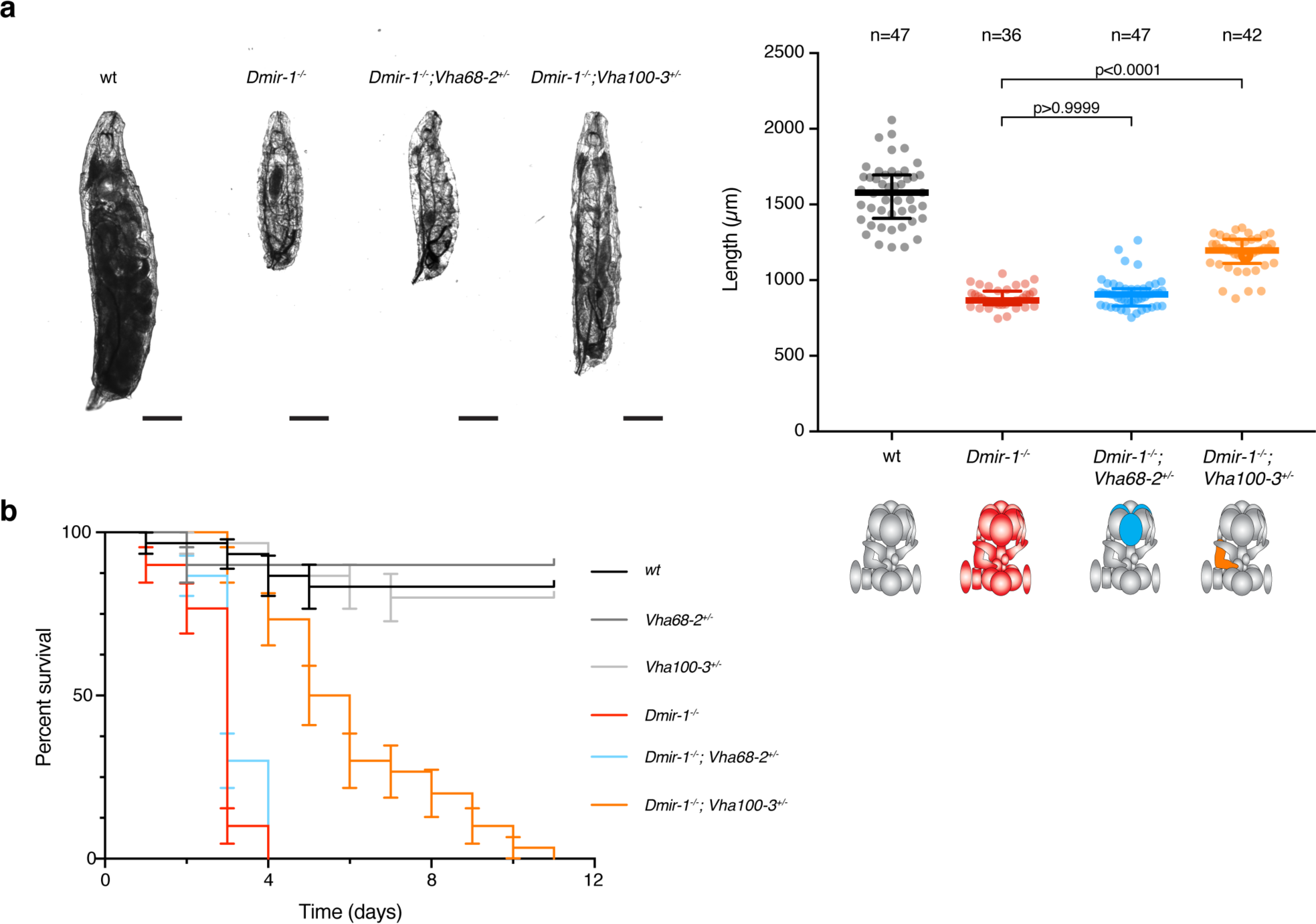
The miR-1/V-ATPase functional interaction is conserved in *Drosophila*. **a**. Reduction in *Vha100-3*, but not *Vha68-2*, copy number, partially rescues *Dmir-1(0)* growth defects. (left) Representative pictures of 24h-post-hatching larvae. Scale bars = 200µm. (right) Quantification of body length. n = number of animals analyzed. P-values by Kruskal-Wallis test are shown. **b**. Reduction in *Vha100-3* copy number extends *Dmir-1(0)* life. Shown are average and s.d. of independent biological triplicates, each with 10 larvae.

The diverse range of cellular defects observed in muscles of animals lacking miR-1 had been so far attributed to numerous different targets^6^, each implicated in a distinct aspect of the phenotype. Here, we identified a conserved set of direct targets that can explain the pleiotropic function of miR-1. The mitochondrial fragmentation, protein aggregation, defects in cell-cell fusion and conductance at the neuromuscular junction observed upon miR-1 loss, can ultimately be related to defects in mitochondrial and lysosome function. Dysfunction of these organelles impacts trafficking and sorting of membrane proteins, autophagy, energy balance and metabolic signaling mainly through the TOR and AMPK pathways^2-4,7^. These cellular processes can in turn be linked to the regulon of functionally-related, yet non-redundant, miR-1 targets comprising the V-ATPase complex and DCT-1/BNIP3 (this study) and TBC-7/TBC1D15^20^. De-regulation of the V-ATPase or TBC-7/TBC1D15 directly affect lysosomal function, whereas the primary effect of DCT-1/BNIP3 de-repression will be on mitochondria. However, given the tight connection between both organelles, defects in one have an impact on the other and result in a global loss of homeostasis of organelle dynamics and metabolic transfer^5^.

The miR-1 regulon comprises genes that are broadly, if not ubiquitously, expressed and that are essential for cell viability. This raises the question of why muscle cells need to specifically repress these genes. We hypothesize that this relates to unique aspects of muscle morphology and physiology. First, muscle has a high energy demand and is thus particularly sensitive to regulators of mitochondrial homeostasis, as well as availability of metabolites that serve as energy sources^50^. Moreover, high levels of respiration generate reactive oxygen species and increased mitochondrial damage, which makes the tight regulation of mitophagy a critical requirement^51^. Finally, muscle cells also display unique morphological characteristics: they are densely packed with myofibers and have a modified ER and plasma membrane to maximize conductance and contraction efficiency while maintaining stability under constant mechanical stress^52^. miR-1-mediated control of mitochondrial and lysosome biology may be necessary to achieve a delicate balance between different homeostatic processes in these highly specialized cells. The type of specific repression of ubiquitous genes mediated by miR-1 may provide an entry point to understand the origin of certain cellular vulnerabilities. For instance, germline mutations in the V-ATPase chaperone VMA21 cause a specific myopathy in humans^53^ but spares most other cell types. Based on our findings, we hypothesize that this may be related to the fact that muscle cells repress V-ATPase subunit expression via miR-1.

miRNAs are repressors of gene expression and as such their biological functions stem from lowering the levels of their targets. Here, we show that miR-1 represses many subunits of a multi-protein complex but this repression of individual components is in fact required to promote function of the complex. Our work reveals a need for miR-1-mediated repression to ensure adequate levels and/or stoichiometry of the different subunits to achieve the correct assembly of the V-ATPase complex. Many V-ATPase subunits are transcriptionally co-regulated by bHLH-Zip transcription factors of the MITF family, mostly acting downstream of the TOR pathway^54^. In *C. elegans*, more than half of *vha* genes occur in operons with shared transcriptional regulation (which notably also include genes involved in mitochondrial proteostasis). Our work suggests that transcriptional regulation alone is not sufficient for establishing conditions for correct V-ATPase assembly in muscle cells, but this requires in addition the post-transcriptional regulation provided by miR-1. The results presented here point to a novel role for miRNAs in coordinating assembly of multi-subunit complexes.

## Acknowledgements

We thank Maria Doitsidou and members of the Cochella lab for critical feedback and discussions, Guy Benian for the UNC-89 antibody, Kathrin Gieseler for sharing the strain KAG420, Julia Riedl for help with MATLAB code, the IMP Biooptics facility and the IMBA fly house for critical input in diverse experiments. We also thank WormBase and FlyBase. *C. elegans* strains provided by the CGC (NIH P40 OD010440) and *Drosophila* stocks from the BDSC (NIH P40 OD018537) were used in this study. This work was supported by grant ERC-StG-337161 (FP7/2007-2013) from the European Research Council, and grants SFB-F43-23 and W-1207-B09 from the Austrian Science Fund, to LC. Funding for electron microscopy was in part provided by the European Union (Interreg RIAT-CZ ATCZ40). Basic research at IMP is supported by Boehringer Ingelheim GmbH.

## Author contributions

PGP and LC designed the experiments, wrote the manuscript and prepared the figures with contributions from other authors, PGP conducted most *C. elegans* experiments, EMS and PGP performed *D. melanogaster* experiments, TL established all imaging data analysis protocols, AS generated *luc108* allele and performed EPGs, TLS generated strains and optimized EPGs, MA generated strains, MB and DH performed EM experiments.

## Materials and Methods

### Strain maintenance

All *Drosophila melanogaster* stocks were maintained under standard conditions at 18°C or 25°C. All *C. elegans* strains were maintained on nematode growth media (NGM) plates seeded with OP50 bacteria at 20°C as previously described^1^, unless indicated otherwise. A complete list of strains used in this study is presented in Supplementary Table 1.

### Generation of transgenic strains

Standard microinjection procedures were used to generate transgenic worms with <u>extra chromosomal arrays</u>^2^. Briefly, DNA was injected as complex arrays in the gonads of N2 young adults. Injection mixes contained 1-5 ng/µL of the plasmid of interest and a similar amount of a co-injection marker (*ttx-3p::mCherry* or *elt-2p::DsRed*), in addition to 100 ng/µL of sonicated genomic DNA from *E. coli* OP50. For VMA21 overexpression, 5 ng/µL of WRM065bD07 fosmid containing *R07E5*.*7/vma21* gene were injected. MLC1955 [*lucEx1114(WRM065bD07, ttx-3p::mCherry)*] and MLC1956 [*lucEx1115(WRM065bD07, ttx-3p::mCherry)*] lines were isolated.

<u>Homology-directed genome editing</u> using Cas9, CRISPR RNA (crRNA) and trans-activating crRNA (tracrRNA) ribonucleoprotein complexes in vitro assembled was performed as previously described^3^. Briefly, purified Cas9 and synthetic RNAs were pre-incubated at 37°C for 15 minutes to enable complex formation and injected into the gonad of young adult worms. Alt-R® CRISPR-Cas9 crRNA targeting the DNA sequences available in Supplementary Table 2 were obtained from Integrated DNA Technologies (IDT). *mir-1(luc108)* was created by deleting 117 bp of the *mir-1* locus, removing the whole sequence coding for *mir-1* hairpin. *vha-1(luc132), vha-8(luc135), vha-11(luc130), vha-13(luc133)* and *vha-14(luc134)* were created by replacing the miR-1 binding site (ACATTCCA) with the restriction site of the enzyme NotI (GCGGCCGC) in their 3’UTRs. *vha-12(luc139)* was created by replacing the three miR-1 binding sites with the restriction site of the enzymes NotI, BamHI (GGATCC) and EcoRI (GAATTC) in its 3’UTR. *dct-1(luc145)* was created by replacing the two miR-1 binding sites with the restriction site of the enzyme NotI in its 3’UTR. *vha-1(luc161)* was created by deleting 454 bp of the *vha-1* locus, removing most of the coding sequence. As this deletion was lethal in homozygosis, this allele was balanced with the *hT2* balancer^4^.

### L1 synchronization

For the phenotypic characterization of *mir-1* mutant animals, 10 to 15 young adults were placed in unseeded 6 cm NGM plates with a bit of OP50 bacteria as food source. After 16 hours at 20°C, L1 larvae were collected and proceed to be imaged. For experiments with starved L1s, 15-20 young adults were bleached using hypochlorite solution (1% NaOCl; 1 M NaOH) in unseeded 6 cm NGM plates. Eggs were then washed three times with M9 buffer (22 mM KH2PO4; 42 mM Na2HPO4; 86 mM NaCl; 1 mM MgSO4) and incubated at 20°C for 16 hours. Starved nutritional state is indicated in the corresponding figure legends.

### Plasmid Construction

All constructs were generated by standard molecular cloning procedures with restriction digest, PCR and Gibson assembly. The coding sequences in the constructs were verified by Sanger sequencing. Plasmid maps and sequences are available upon request.

### RT-qPCR

mRNA quantification of *mir-1* predicted target genes in *C. elegans* was assessed via RT-qPCR following the procedure described in^5^. Briefly, 10 starved and synchronized L1s were transferred to 2 μl of Lysis Buffer (5mM Tris-HCl pH 8.0, 0.25 mM EDTA and 1 mg/mL Proteinase K, 0.5% Triton X-100, 0.5% Tween20) using an eyelash. These samples were subjected to 10 minutes digestion at 65°C before heat inactivation of proteinase K for 1 min at 85°C. Genomic DNA was removed by incubating the samples with DNaseI (NEB, cat #M0303S) for 10 minutes at 37°C. Then, DNaseI was inactivated by adding 25 mM EDTA and incubating the samples for 10 minutes at 65°C. Next, crude lysates were reverse transcribed (Thermo Fisher, cat #4368814) before performing qPCR using the GoTAQ qPCR Mastermix (Promega, cat #A6001) according to the manufacturer’s instructions. Relative expression was calculated according to the ΔΔCq-method using *cdc-42* as a reference gene.

For mRNA quantification of *D. melanogaster vha* subunits, 100 newly hatched L1s of the indicated genotype were transferred to 500 µl Trizol reagent (Thermo Fisher, cat. #15596026) and homogeneized using an electric pestle tissue grinder for 2-3 minutes. Total RNA was isolated following the manufacturer’s instructions, and genomic DNA was removed by incubating with DNAse I (NEB, cat #M0303S). Samples (1 µg total RNA) where then reverse-transcribed using Superscript IV Reverse Transcriptase (Thermo-Fisher, cat. #18090010), with oligo-dT as a primer. The resulting cDNA was used for qPCR, using the the GoTAQ qPCR Mastermix (Promega, cat #A6001) according to the manufacturer’s instructions. For each sample, three technical replicates and an RT-control were performed. The standard deviation between technical replicates was always smaller than 0.5. In each reaction, a cDNA standard sample was included and its Cq value was checked not to differ by more than 0.5 compared to the same standard sample used in the calibration curve of each gene. Relative expression was calculated according to the ΔΔCq-method, using *actin42a* and *ef1α2* as reference genes. A complete list of primer sequences is provided in Supplementary Table 3.

### Cryo-preparation of *C. elegans*

N2 and MLC1384 [*mir-1(luc108) I*] animals were cultivated on three 15 cm peptone enriched plates seeded with concentrated HB101 *E. coli* bacteria. Adult worms were bleached using hypochlorite solution and isolated eggs were washed three times in M9 buffer, as previously described^6^. After approximately 16 hours, a pellet of L1 worms, containing 5% BSA (Fraction 5) in M9 buffer was transferred into the 100 µm cavity of a 3 mm aluminum specimen carrier. This carrier was sandwiched with a flat 3 mm aluminum carrier and immediately frozen under high pressure in a HPF Compact 01 (Engineering Office M. Wohlwend GmbH). The frozen samples were subsequently transferred into a Leica EM AFS-2 freeze substitution unit (Leica Microsystems). Over a period of five days, samples were substituted in a medium of acetone containing 2% osmium tetroxide (EMS), 0.2% uranyl acetate (Merck) and 5% water. Freeze substitution was performed according to the following protocol: 60 hours at –90°C, warm up at a rate of 2°C per hour to –54°C, 8 hours at –54°C, warm up at a rate of 5°C per hour to –24, 15 hours at -24°C, warm up at a rate of 6°C per hour to 20°C, 5 hours at 20°C. At 20°C samples were taken out and washed 3 times in anhydrous acetone. To enhance contrast the samples were stained using 1% thiocarbohydrazid (Sigma Aldrich) for 20 minutes at RT, 2% osmium tetroxide for 20 minutes at room temperature and 2% uranyl acetate for 20 minutes at 60°C (three times 10 minutes washing steps after each staining step). Next, samples were infiltrated with Agar 100 Epoxy resin (Agar Scientific), in a graded series of acetone and resin over a period of 2-3 days. Polymerization took place at 60°C.

Ultra-thin sections were cut using a Leica UCT ultramicrotome (Leica Microsystems) at a nominal thickness of 70 nm and picked up on 100mesh Cu/Pd grids (Agar Scientific) previously coated with a supporting film of formvar (Agar Scientific). Examination regions on the sections were selected at random, inspected with a FEI Morgagni 268D (FEI, The Netherlands) operated at 80 kV. Digital images were acquired using a 11-megapixel Morada CCD camera (Olympus-SIS, Germany).

### Behavior and morphological assays

#### Mitochondria network

For analysis of mitochondrial network organization in C. elegans body-wall muscles, the strain SJ4103 [*zcIs14(myo-3::GFP(mit))*] was used^7^. Images from synchronized L1s were deconvoluted with CMLE algorithm, 40 iterations and 15 Signal/Noise ratio (Huygens deconvolution software, Scientific Volume Imaging) and quantified using a two-step approach: We applied a 3D Laplacian of Gaussian filter and thresholded the resulting image (Fiji-ImageJ software, version 1.52p). This binary image was used to create surface objects in Imaris software (version 9.3.1, Bitplane), allowing for measurement of their area, volume and sphericity and export the values for statistical analysis. An average of 150-200 mitochondria objects (n) was detected per animal (N). Images with less than 50 objects detected were excluded for further analysis.

#### ATP levels

For measuring ATP levels in the body-wall muscle of C. elegans, the strain GA2001 [*wuIs305 (myo-3p::Queen-2m)*] expressing the ratiometric Queen-2m ATP biosensor was used as previously described^8^. Images from fed and synchronized L1s were further processed in Fiji-ImageJ software. After subtracting the background, an average intensity projection of the whole Z-stack was applied for both 405nm and 488nm excitation channels. Then, the ratio 405ex/488ex was calculated by dividing the 32-bit resulting images. Three independent regions of interests (ROIs) were selected and the median of them was used for plotting and statistical analysis.

#### Protein aggregation

To assess proteostasis defects, synchronized and fed L4s animals carrying the rmIs133 [*unc-54p::Q40::YFP*] transgene in their body-wall muscles were imaged^9^. The resulting images were further processed with Fiji-ImageJ software: a maximum-intensity Z-stack projection was extracted followed by thresholding. Particles with a circularity between 0.5-1.0 and size bigger than 4 pixels were counted as aggregates.

#### pm3D cell-cell fusion

pm3DL and R fusion was assessed by the expression of a pm3DR-specific membrane marker (*mir-4813p::myr::GFP*). Images from synchronized and fed L1s were deconvoluted with CMLE algorithm, 40 iterations and 15 Signal/Noise ratio (Huygens deconvolution software, Scientific Volume Imaging). To obtain representative cross sections of pm3D, we used an ImageJ-Macro which allowed us to automatically export three equidistant cross-sections perpendicular to a manually drawn line along the respective region. For the export the “Reslice”-function was used. Cross-sections were analyzed blindly. Macro is available upon request.

#### Recording Electropharyngeograms (EPGs)

EPGs were used as a read-out of pharyngeal pumping in N2, MT17810 [*mir-1(n4102) I*] and MLC1384 [*mir-1(luc108) I*]. Approximately 22 hours before recording, about 40 L4s were transferred onto an NGM plate seeded with OP50 at 20°C. Prior to starting the recordings, the worms were washed with 1.2ml M9 to remove pieces of agar and food. After removing 1ml of the supernatant, 1ml of 10mM serotonin (diluted in M9) was added to the worms. Worms were incubated in serotonin for 40 minutes before loading them on the ScreenChip^™^ System (SC40, InVivo Biosystems, #cat. SKC101). For acquiring EPG-data as well as data analysis we used the software supplied by InVivo Biosystems (NemAquire and NemAnalysis). Pharyngeal pumping of each worm was recorded for two minutes. All recordings with a signal-to-noise ratio level below 3.0 were discarded. For comparing the peak-triggered average between different worm strains, we used MATLAB in collaboration with Julia Riedl (Zimmer lab, IMP).

#### Drosophila larval survival assay

Adult flies were allowed to lay embryos on apple juice agar plates for 4 h at 25 °C. On the next day, 10 newly hatched L1s of the desired genotype were collected and placed on fresh apple juice agar plates with food (paste of baker’s yeast and water). Every 24 hours, alive larvae were transferred to fresh plates with food, while dead larvae were removed and scored. The experiment was followed until all of the larva either turned into a pupa or died.

#### Drosophila larval length

Adult flies were allowed to lay embryos on apple juice agar plates for 4 hours at 25°C. On the next day, 10 newly hatched L1s of the desired genotype were collected and placed on fresh apple juice agar plates with food (paste of baker’s yeast and water). After 24 hours, the larvae were collected and anesthetized in a small etherization cage using a 1.5 mL microcentrifuge tube, as previously described^10,11^. Larvae were then mounted in a slide with a 2.5% bacto-agar pad and imaged in a widefield upright microscope (Zeiss Axio Imager.Z2 with sCMOS camera, 5x objective). Length of the larvae was measured using Fiji-ImageJ software.

### Spinning Disk Confocal Microscopy

L1 or L4 worms were mounted on a 5% bacto-agar pad and immobilised in 50mM NaN3. Images were acquired using a Visiscope Spinning Disc Confocal (Visitron Systems GmbH, Puching, Germany) with PCO Edge 4.2m sCMOS camera, CFI plan Apo lambda 100x/1.45 oil (or 10x/0.45 air for protein aggregation) and 100% power of lasers 405nm 120mW, 488 nm 200 mW and/or 561 nm 150 mW. Z-stack space 0.1 µm, except for the analysis of protein aggregation that a Z-stack space of 1 µm was used.

### Data analysis

Statistical tests were performed in Prism v7 or 8 (GraphPad Software Inc.) and are described for each figure. Generally, to identify outliers, the ROUT test was applied. To assess if the data was following a normal distribution, three independent normality tests (D’Agostino and Pearson, Shapiro-Wilk and KS) were performed. If the distribution was normal, One-way analysis of variance (ANOVA) was performed for comparisons across multiple independent samples, using Tukey’s multiple comparisons correction. If not, Kruskal-Wallis test using Dunn’s multiple comparisons correction was used. Median with interquartile range was plotted for all behavior analysis. For RT-qPCR analysis, an unpaired t-test was performed.

## Extended Data Figures

**Extended data Figure 1.**
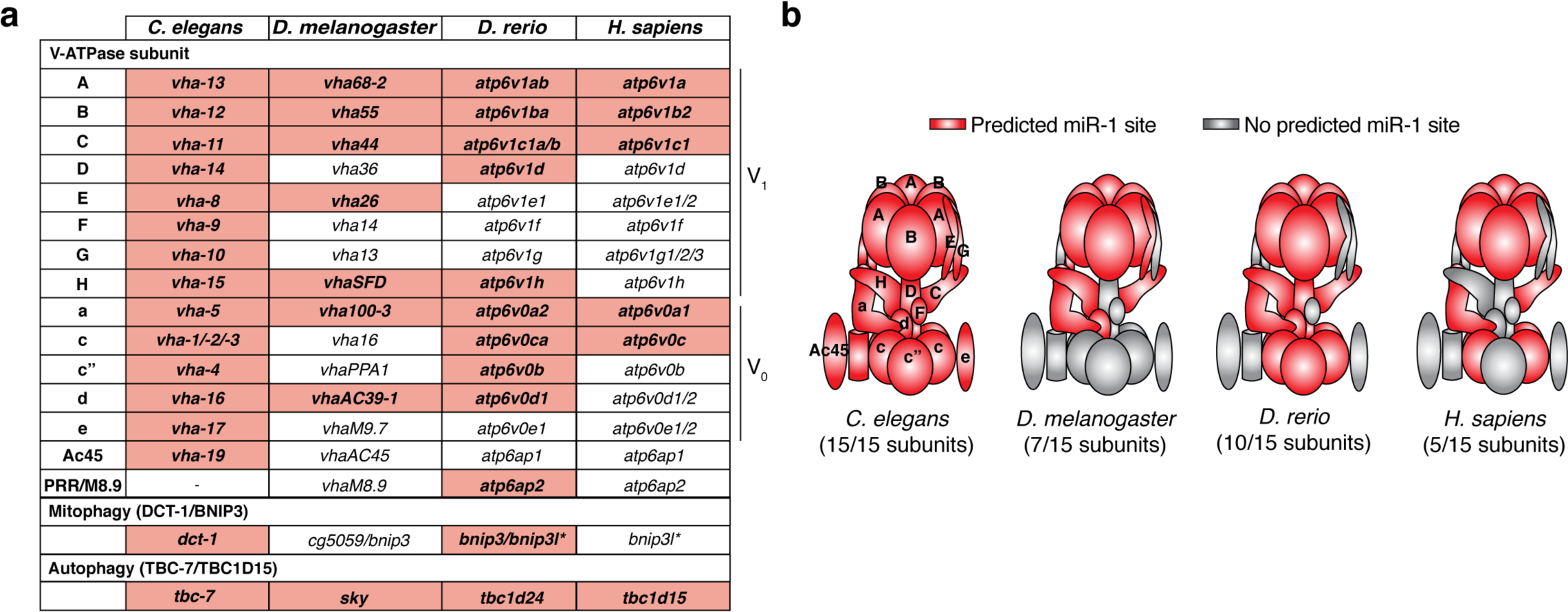
V-ATPase, DCT-1/BNIP3 and TBC-7/TBC1D15 are predicted miR-1 targets across animals. **a**. Several subunits of the V-ATPase complex, *dct-1/bnip3* and *tbc-7/tbc1d15* have at least one miR-1 predicted binding site in their 3’ UTR (bold, red shade) in *C. elegans, D. melanogaster, D. rerio* and *H. sapiens*^12-14^. Asterisks (*) refer to the presence of a predicted miR-133 binding site in *bnip3l* instead; in vertebrates, miR-1 and miR-133 are clustered together and co-expressed specifically in muscle^15^. **b**. Location of V-ATPase subunits that are predicted miR-1 targets; miR-1 regulation could affect subunits across the whole complex.

**Extended Data Figure 2.**
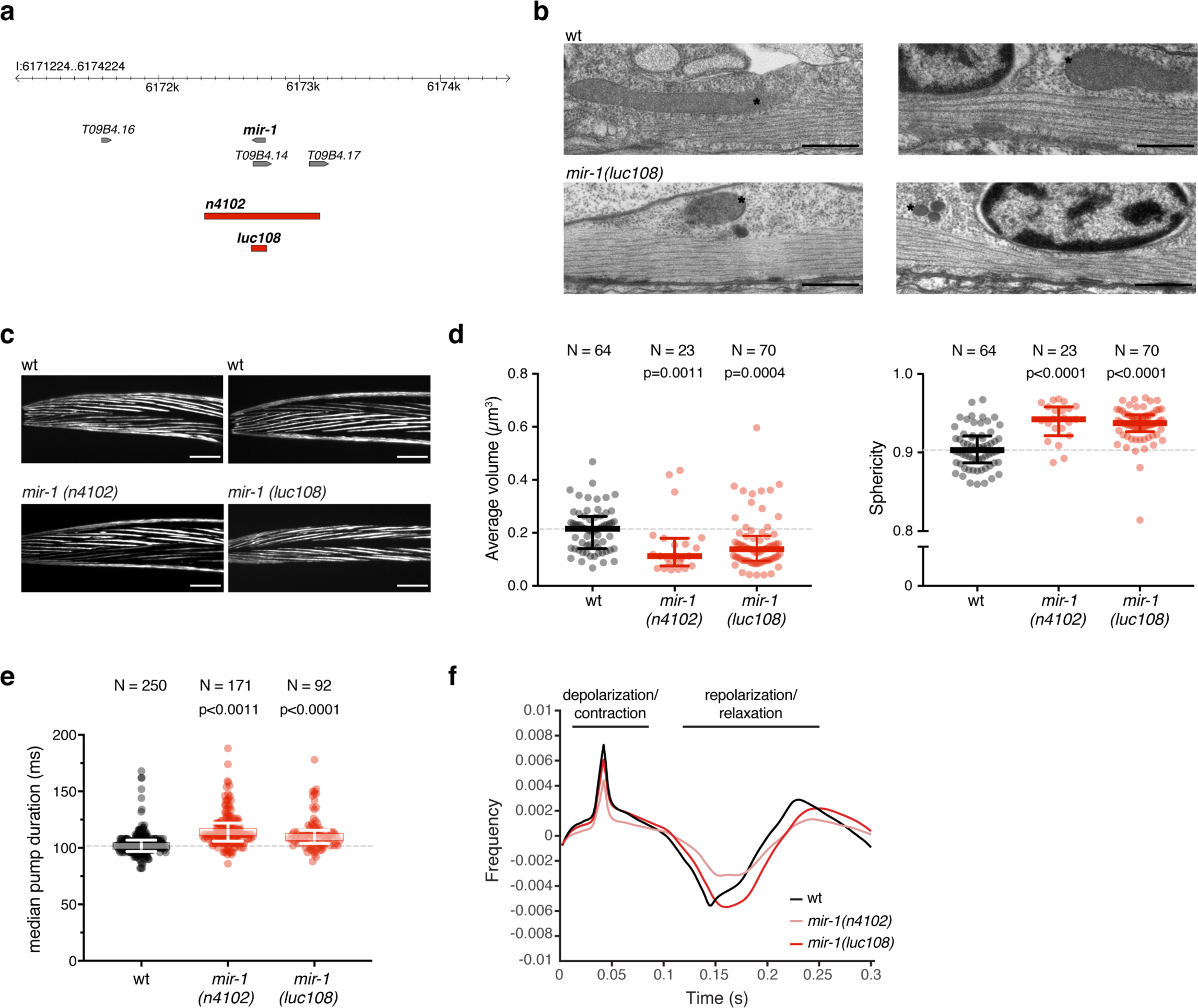
Absence of miR-1 causes diverse muscle defects in *C. elegans*. **a**. A schematic depicting the *mir-1* locus and the two deletion alleles used in this study: *n4102* was obtained by EMS mutagenesis^16^, *luc108* was generated using CRISPR/Cas9 for this study. **b**. Representative electron microscopy images for both wt and *mir-1(luc108)* L1 larvae. The sarcomere structure is preserved, asterisks indicate mitochondria. Scale bars = 0.5µm. **c**. Representative fluorescence images of animals expression MYO-3::GFP (maximum intensity projections of stacks through the whole animal). The typical parallel organization is preserved in both *mir-1(0)* L1s alleles. Scale bars = 10µm. **d**. (left) Average mitochondria volume, extracted from confocal microscopy images of L1 larvae expressing mitochondrial GFP (see Fig. 1a and Methods). (right) Mitochondria sphericity was extracted from the same images. Mitochondria are smaller and more spherical in *mir-1(0)* animals, indicating fragmentation of the network. P-values by Kruskal-Wallis test are indicated. **e**. Median duration of each pharyngeal pumping cycle was extracted from the EPGs analyzed in Fig. 1e using NemAcquire and NemAnalysis softwares. In *mir-1(0)* animals each pump is longer, causing the lower overall frequency (Fig. 1e). P-values by Kruskal-Wallis test are indicated. **f**. Peak-triggered average of all pharyngeal pumping events for each genotype revealed a normal duration of the contraction phase, but a longer relaxation/recovery phase in *mir-1(0)* young adults, suggesting a reduced capacity of recovering after each contraction. N/n refers to number of animals analyzed.

**Extended Data Figure 3.**
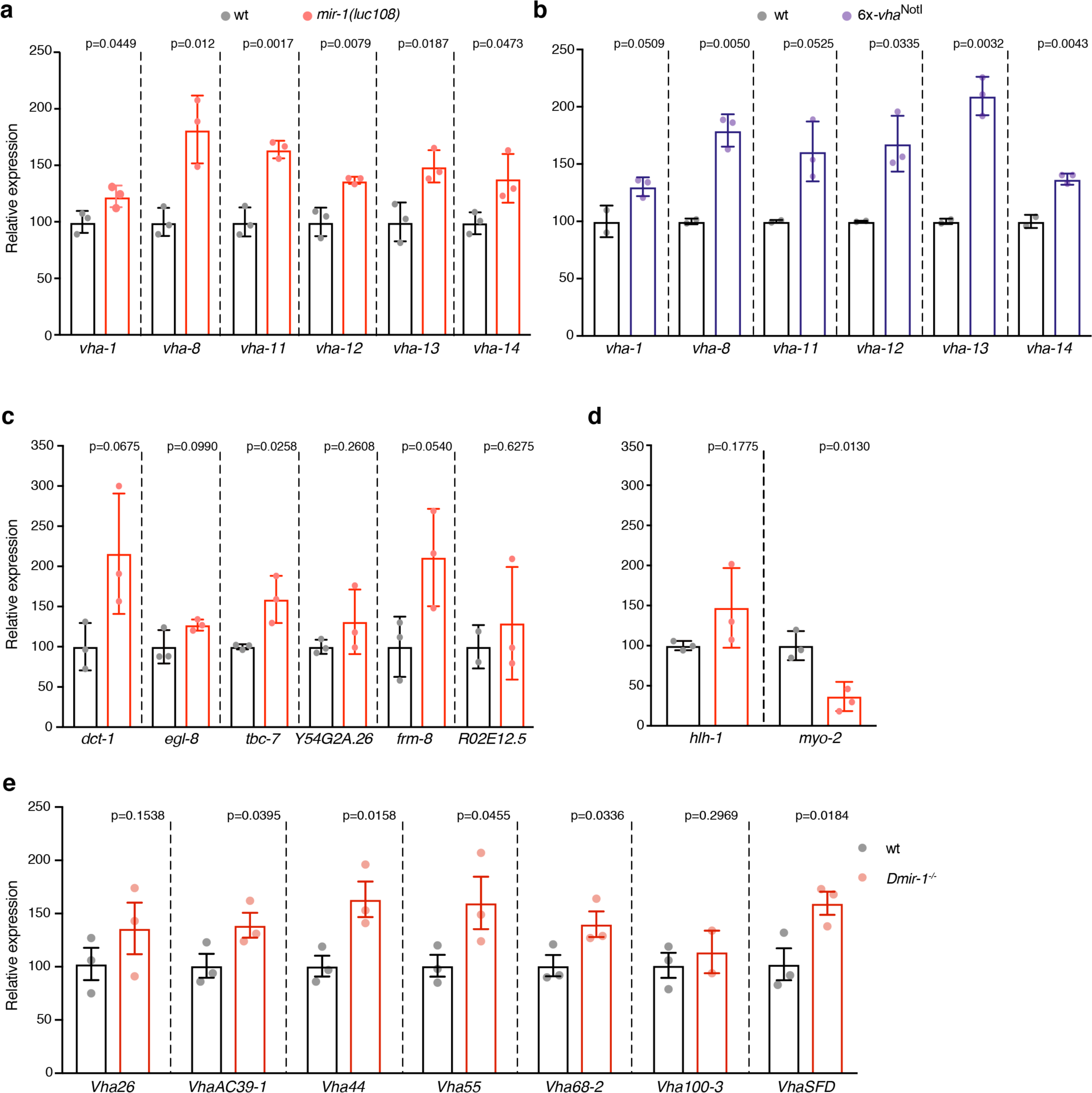
Multiple subunits of the V-ATPase complex are repressed by miR-1 in *C. elegans* and *D. melanogaster*. **a-d**. RT-qPCR on whole L1 larvae of the indicated genotypes for a number of predicted miR-1 targets (**a-c**) and controls (**d**). *vha-1, -8, -11, -12, -13* and *-14* mRNA levels are increased in both *mir-1(0)* and *6x-vha*^*NotI*^ mutants, indicating they are direct targets of miR-1. **c**. *dct-1* and *tbc-7* are significantly upregulated in *mir-1(0)* animals. **d**. *mir-1(0)* animals show defects in muscle differentiation, as shown by increased levels of the body wall muscle-master regulator (*hlh-1*) and decreased levels of pharyngeal myosin (*myo-2*). Three independent biological replicates were analyzed, except for the wt samples in panel b (two biological replicates tested). Each biological replicate consists of 10 starved L1 animals. **e**. RT-qPCR for various *Vha* transcripts from whole *Drosophila* first instar larvae of the indicated genotypes; *Vha* transcripts are upregulated in the absence of miR-1. Three independent biological replicates, consisting of 10 larvae, were analyzed. P-values by t-test are shown.

**Table S1.**
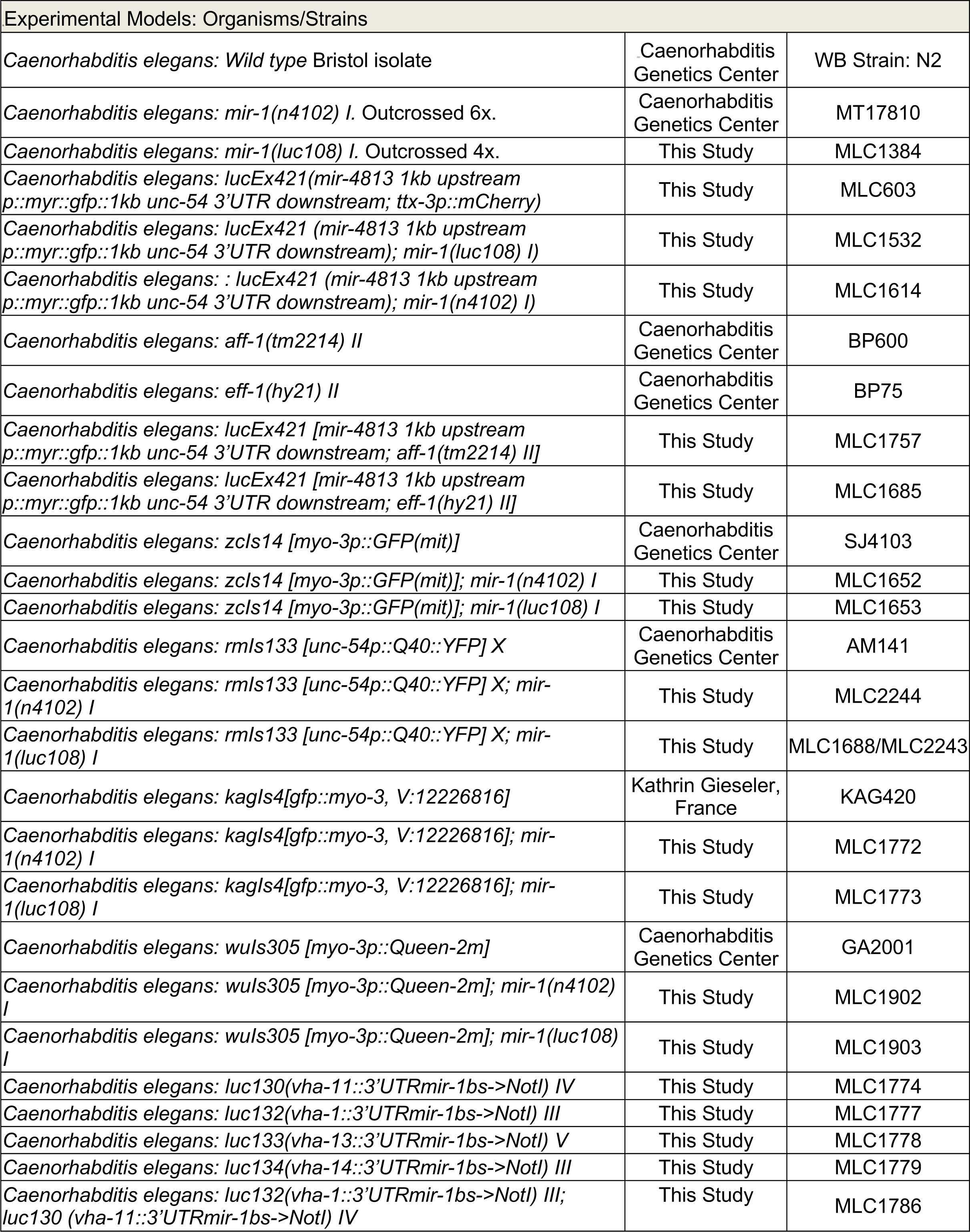
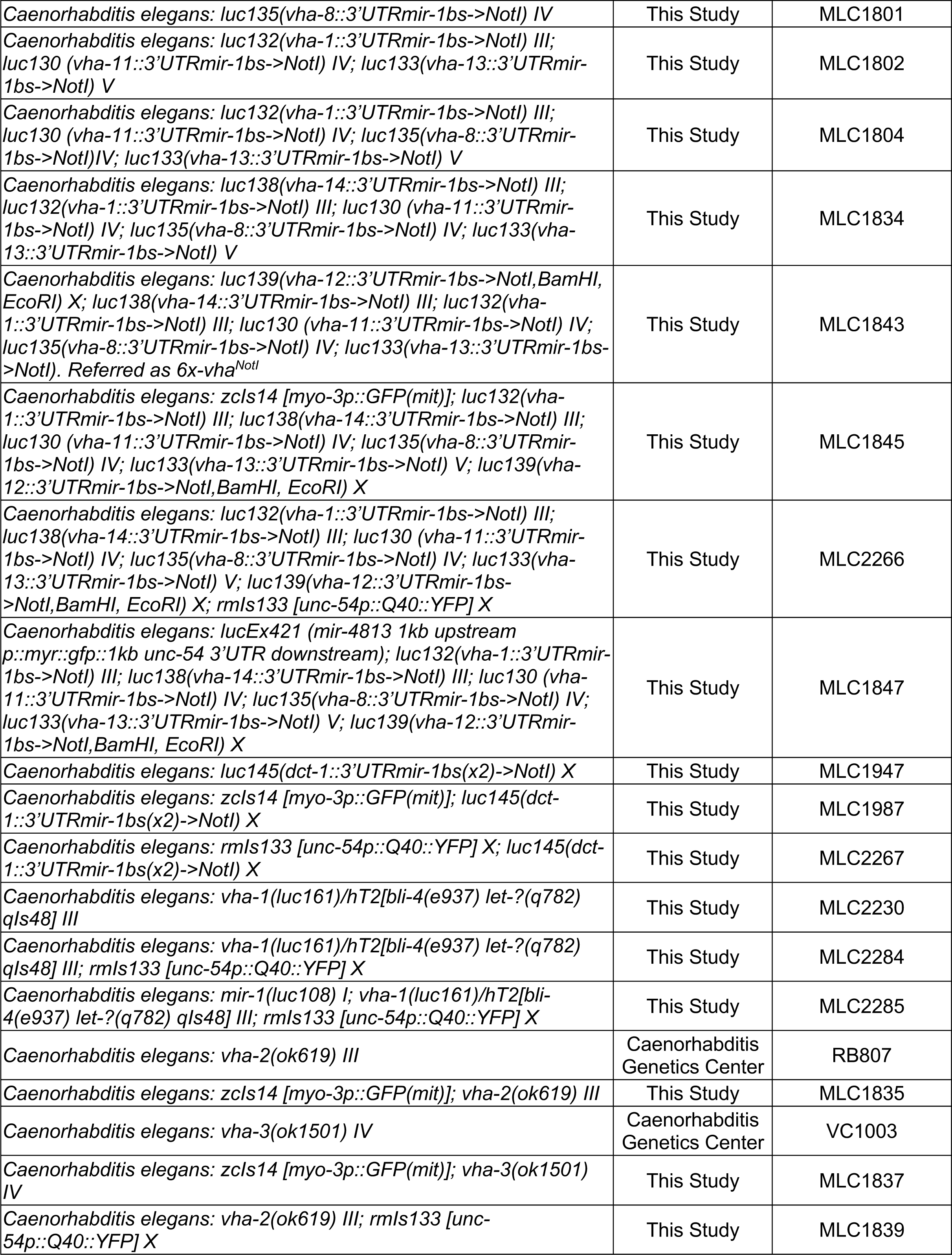
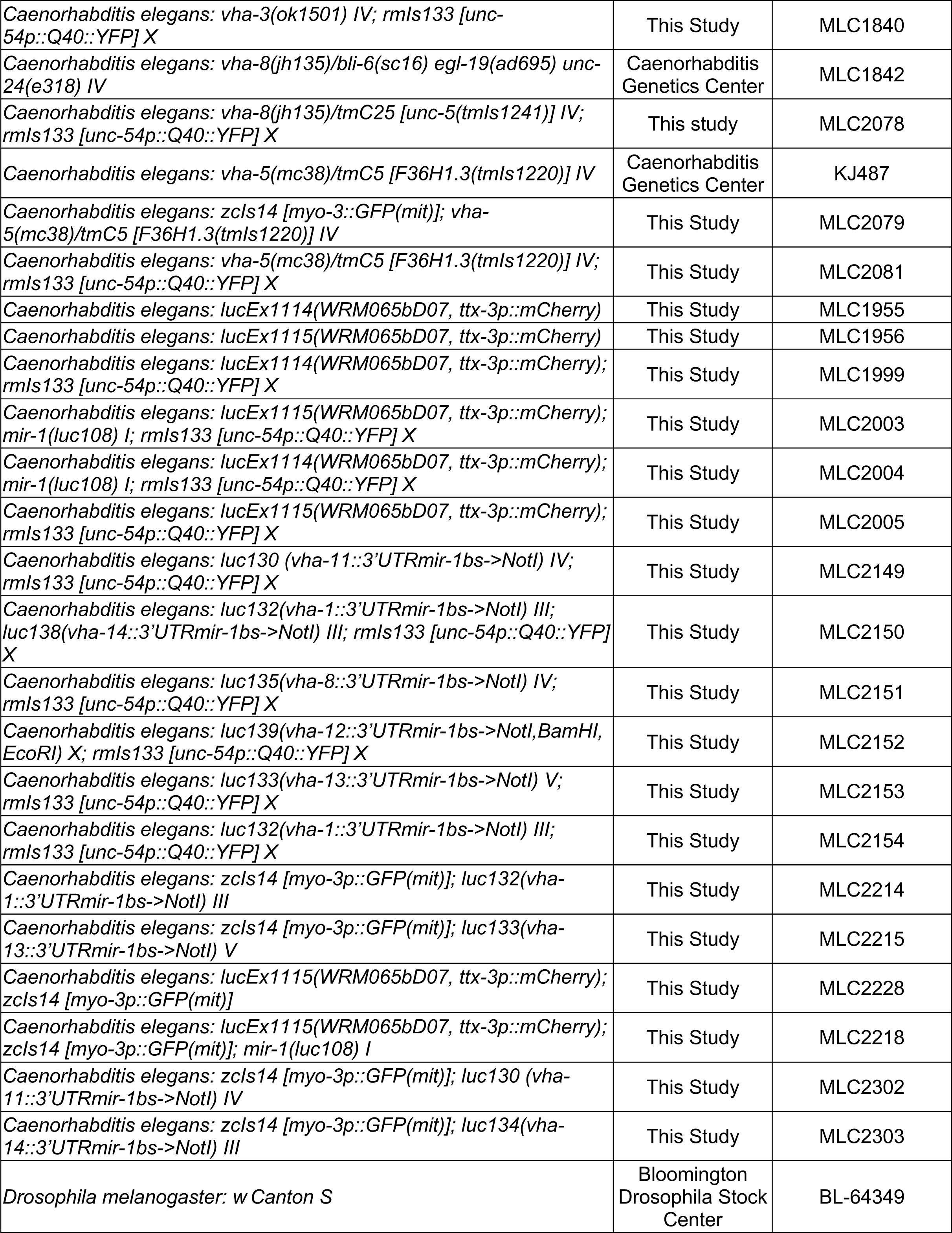
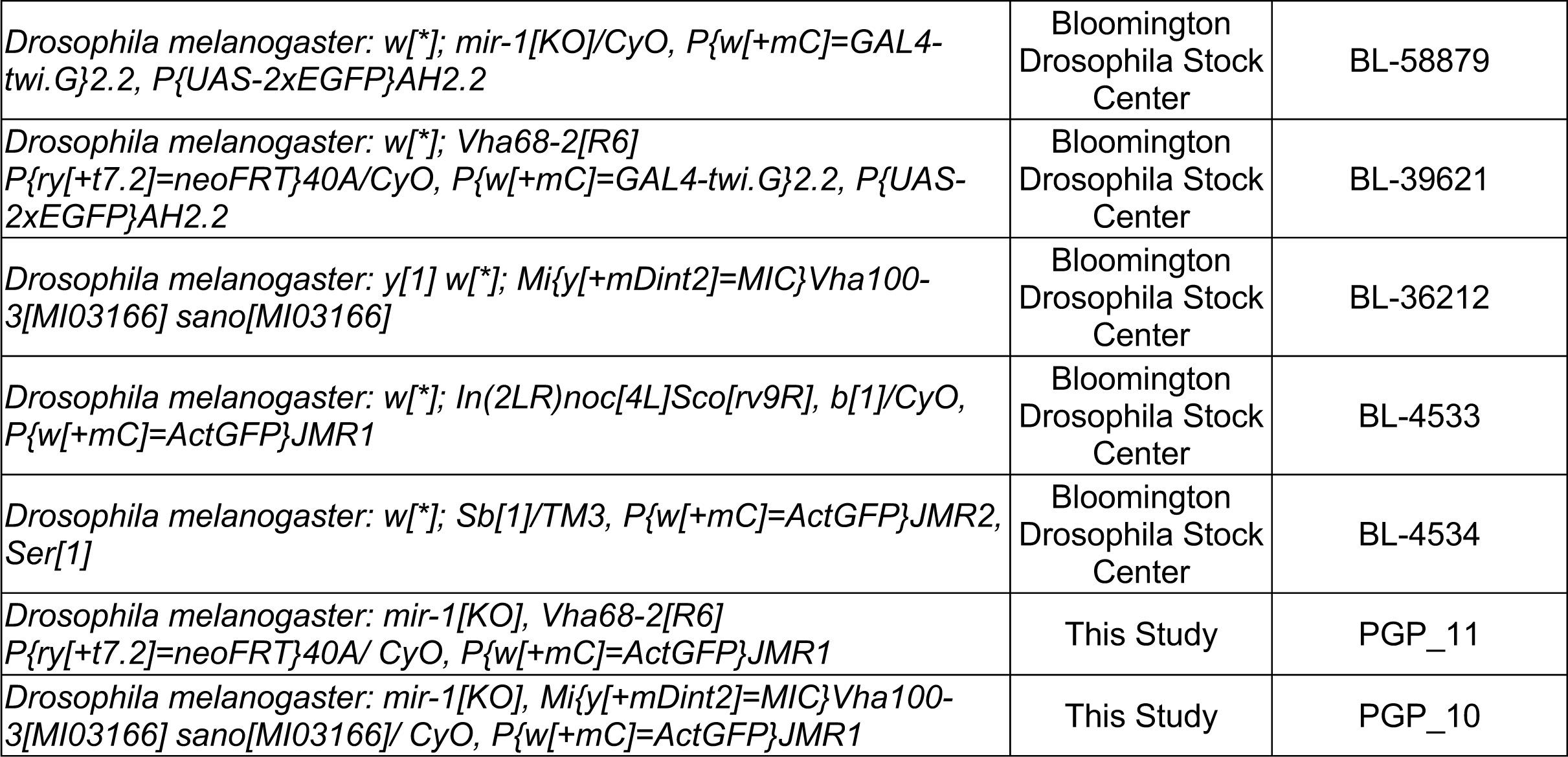
Experiment Models: Organisms/Strains used in this study.

**Table S2.**
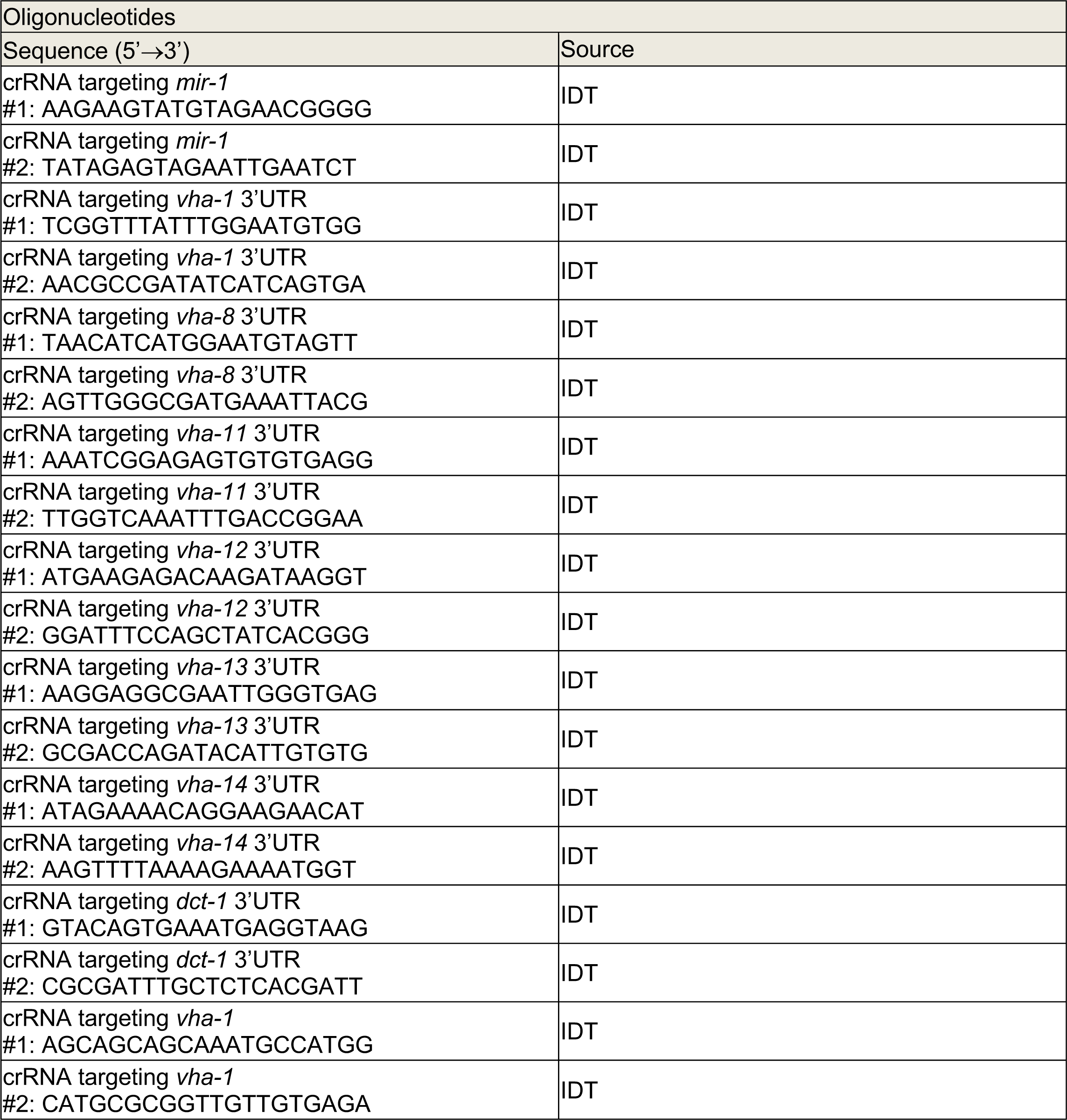
Alt-R® CRISPR-Cas9 crRNA sequences for Homology-directed genome editing used in this study.

**Table S3.**
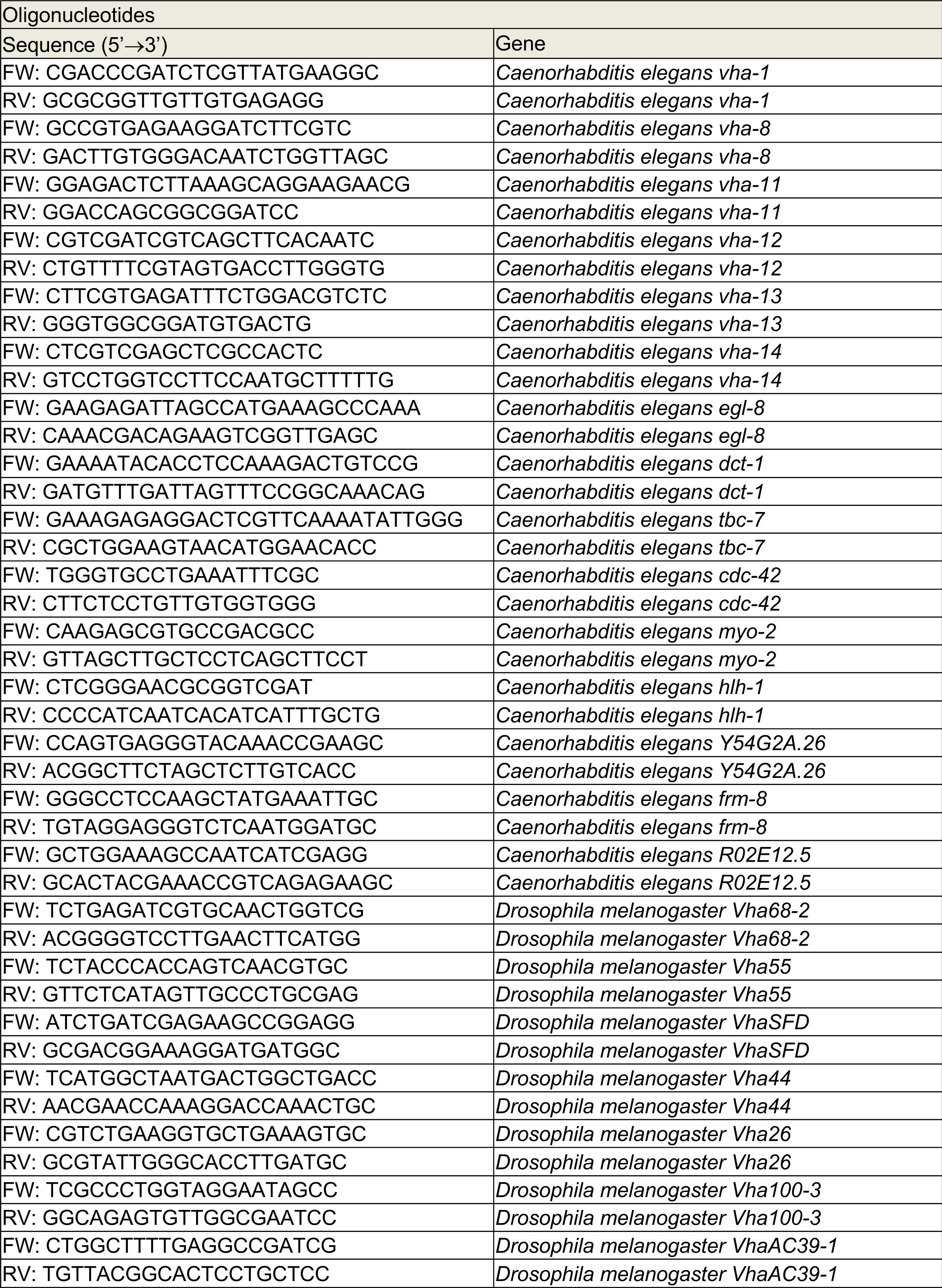
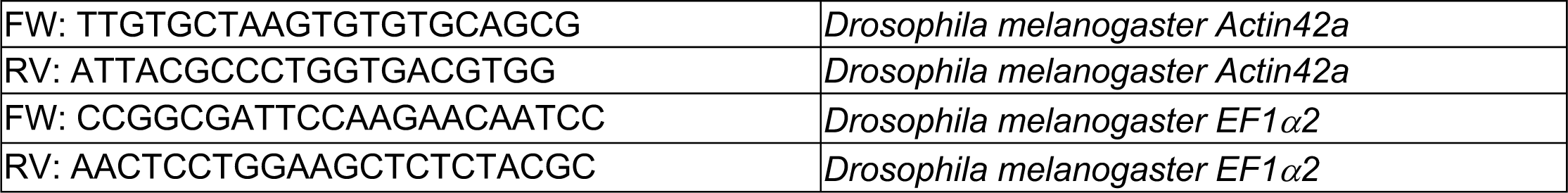
Oligonucleotides sequences for qPCR used in this study.

